# Reconstitution of contractile actomyosin rings in vesicles

**DOI:** 10.1101/2020.06.30.180901

**Authors:** Thomas Litschel, Charlotte F. Kelley, Danielle Holz, Maral Adeli Koudehi, Sven Kenjiro Vogel, Laura Burbaum, Naoko Mizuno, Dimitrios Vavylonis, Petra Schwille

## Abstract

One of the grand challenges of bottom-up synthetic biology is the development of minimal machineries for cell division. The mechanical transformation of large-scale compartments, such as Giant Unilamellar Vesicles (GUVs), requires the geometry-specific coordination of active elements, several orders of magnitude larger than the molecular scale. Of all cytoskeletal structures, large-scale actomyosin rings appear to be the most promising cellular elements to accomplish this task. Here, we have adopted advanced encapsulation methods to study bundled actin filaments in GUVs and compare our results with theoretical modeling. By changing few key parameters, actin polymerization can be differentiated to resemble various types of networks in living cells. Importantly, we find membrane binding to be crucial for the robust condensation into a single actin ring in spherical vesicles, as predicted by theoretical considerations. Upon force generation by ATP-driven myosin motors, these ring-like actin structures contract and locally constrict the vesicle, forming furrow-like deformations. On the other hand, cortex-like actin networks are shown to induce and stabilize deformations from spherical shapes.

## Introduction

In cells, actin filaments are organized into cross-linked, branched, and bundled networks. These different architectures appear in structures such as filopodia, stress fibers, the cell cortex, and contractile actomyosin rings; each has unique physical properties and fulfills different roles in important cellular processes.^1^ These different structures must be actively assembled and maintained by cellular factors, such as the many actin cross-linking proteins. By mediating higher-order actin organization, cross-linkers allow actin filaments to fill a diverse array of structural and functional roles within cells.^2,3^

In many cases, actin networks are linked to, or organized around, cellular membranes. Actin polymerization is a driving force behind many examples of membrane dynamics, including cell motility, membrane trafficking, and cell division.^1^ Many of the actin binding proteins involved in these processes are directly regulated via interactions with phospholipid bilayers^4,5^ and membrane interactions have, in turn, been shown to physically guide actin assembly.^6^ While the link between the actin cytoskeleton and phospholipid bilayers is clear, how these connections affect the large scale organization of complex actin networks remains an open question.

Actin is not only one of the most prevalent proteins in current reconstitution experiments,^7,8^ but also was one of the first proteins to be explored in such approaches.^9,10^ The focus of actin related work has since shifted from identifying the components responsible for muscle contraction,^11^ to investigating more detailed aspects of the cytoskeleton^8,12^, such as the dynamics of actin assembly^13,14^ or the cross-talk with other cytoskeletal elements.^15^ These experiments have extended to actin-membrane interactions, including reconstitution of actin cortices on the outside of giant unilamellar vesicles (GUVs),^16-18^ and contractile actomyosin networks associated with supported membranes.^19-21^ Recently, creating a synthetic cell with minimal components recapitulating crucial life processes, such as self-organization, homeostasis, and replication, has become an attractive goal.^22,23^ As such, there is increased interest in work with actin in confinement and specifically within GUVs^24,25^, in order to mimic cellular mechanics, by encapsulating actin and actin binding proteins in vesicles.^26-28^ However, the investigation of higher-order actin structures or networks has been the subject of few studies thus far.^28-31^

Due to the difficulty of encapsulating functional proteins within membrane vesicles, much of the past work has been limited to adding proteins to the outside of vesicles or onto supported lipid membrane systems. However, novel encapsulation methods such as cDICE, as used here, have enabled the efficient transfer of proteins and other biomolecules into cell-sized phospholipid vesicles, as an ideal setting to study complex cellular processes involving membranes.^32-35^ Here, we optimized the encapsulation procedure for a high degree of reproducibility and precision. This did not only allow us to reconstitute novel cell-like cytoskeletal features, but helped to achieve a high level of consistency and efficiency required for comparison of our experimental results with numerical simulations of confined interacting actin filaments. The development of experimentally testable predictive theoretical models is central for the future design of complex experiments that approach the functional complexity of biological systems.

We combined actin bundling and actin-membrane linkage to obtain results more closely resembling *in vivo* morphologies than previously achieved *in vitro*. Specifically, we induced the formation of membrane-bound single actin rings, which imitate the contractile division rings observed in many cells. In agreement with our numerical simulations, we show that membrane anchoring significantly promotes the formation of actin rings inside vesicles. We achieved close to 100% probability of ring formation in vesicles when using the focal adhesion proteins talin and vinculin, which we recently identified as effective actin bundlers [Kelley et al., in revision]. With the inclusion of motor proteins, these actomyosin rings contract similar to those observed in yeast protoplasts.^36^ This work brings us much closer to our goal of being able to quantitatively design and experimentally achieve full division of a synthetic membrane compartment, and thus, to the self-reproduction of artificial cells, a persistent goal in bottom-up biology.^37-40^

## Experimental System

In order to investigate the interplay between actin cross-linking and membrane binding, we used a modified continuous droplet interface crossing encapsulation method (cDICE)^41,42^ to encapsulate G-actin with associated proteins and generate cytoskeletal giant unilamellar vesicles (GUVs) made from the lipid POPC (Figure 1A). Since components cannot be added once the reaction mix is encapsulated, the precise composition of the initial reaction mix is crucial. By tuning concentrations of the polymerization buffer, bundling proteins, membrane anchors and motor proteins, we manipulated the final morphology of the actin network.

**Figure 1:**
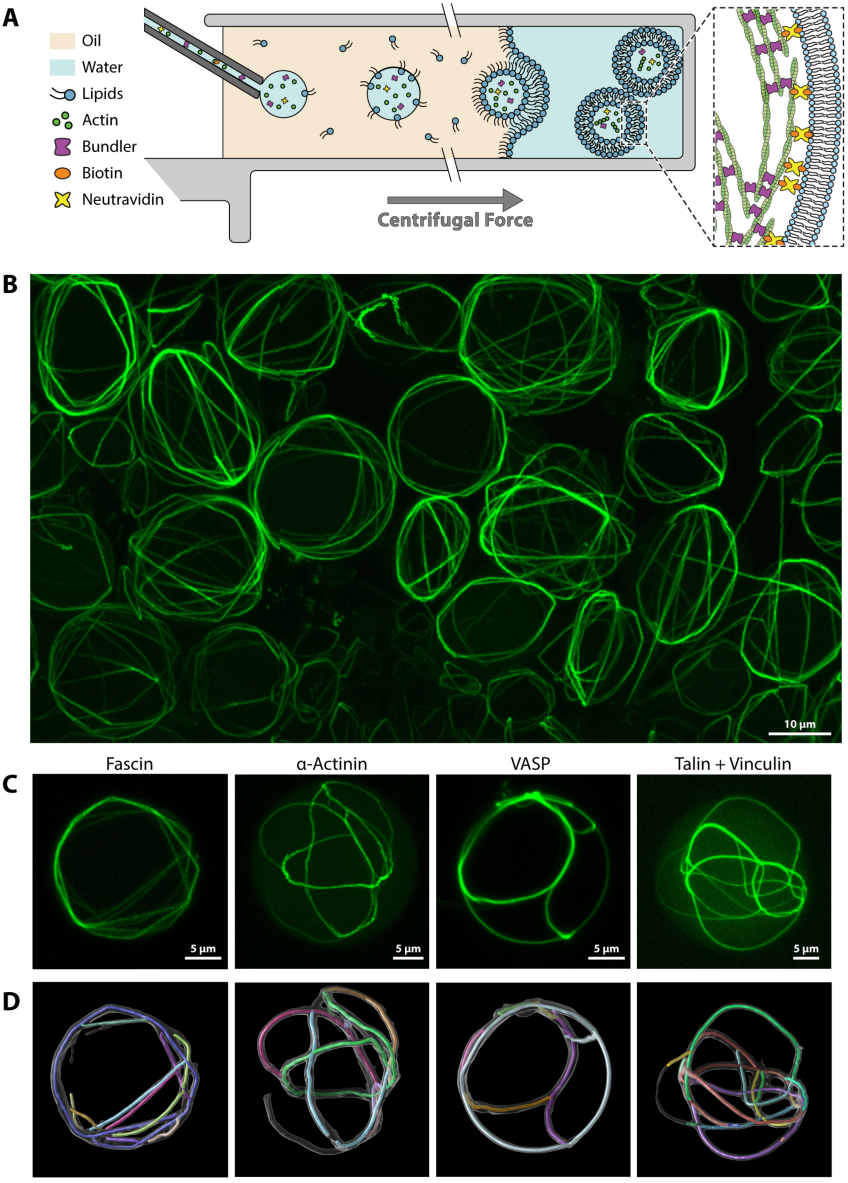
Encapsulation of bundled actin in giant unilamellar vesicles. **A)** Schematic depiction of the vesicle generation process. The aqueous protein solution is injected into a rotating chamber through a glass capillary. Droplets form at the capillary tip in the oil phase, which contains lipids. The droplets then pass through a water–oil interphase lined with a second lipid monolayer, forming the GUVs. **B)** Field of view image (Z-projections of confocal stacks) with many cytoskeletal vesicles. Actin in green. See Movie S1 for 3D effect. **C)** Comparison of cytoskeletal vesicles with actin bundled by 4 different types of bundling proteins. Supplementary Movie S2 shows a 3D view of these vesicles. We used 2 µM actin in all cases, but due to differences in bundling activity, different concentrations of bundling protein: 0.3 µM fascin, 0.9 µM VASP, 1 µM α-actinin, 2 µM talin and 2 µM vinculin. **D)** Automated tracing of bundles by analysis script. Confocal z-stacks are converted into a 3-dimensional “skeleton”-model.

By co-encapsulating actin with known actin cross-linking proteins, we achieved large-scale networks with clearly discernible actin structures. To determine whether different bundling proteins could achieve unique higher-order actin networks, we tested four different types of actin bundling proteins: Fascin, α-actinin, vasodilator-stimulated phosphoprotein (VASP) and a combination of the focal adhesion proteins talin and vinculin. Each case represents a slightly different mechanism of actin binding. Fascin, a 55 kDa protein, binds to actin through two distinct actin binding sites, thereby inducing filament cross-links as a monomer.^43^ α-Actinin, 110 kDa, forms a dimer which bridges two filaments^29,44^. Talin (272 kDa) and vinculin (116 kDa) both dimerize and also require interactions with each other in order to bind and bundle actin filaments [Kelley et al., in revision]. Here we use a deregulated vinculin mutant (see supplementary information). VASP (50 kDa) forms a tetramer, which can link up to four filaments together.^45^ Under all four conditions, the formation of thick filament bundles was observed (Figure 1B,C). Interestingly, while α-actinin, talin/vinculin, and VASP all produced similar morphologies, fascin bundles take on the most unique appearance. These bundles often bend in kinks when their path is obstructed by the membrane, while the other proteins form smoothly curved bundles that can follow the curvature of the encapsulating membrane (Figure 1C,D).

After establishing successful encapsulation of actin and its bundling proteins, we modified the approach and linked the actin filaments to the phospholipid bilayer via biotin-neutravidin bonds, similar to previous work on planar supported lipid bilayers.^19^ This requires the incorporation of biotinylated lipids in the vesicle membranes and the addition of both, biotinylated g-actin and neutravidin in the encapsulated reaction mix. We tested different fractions of biotinylated lipids as well as biotinylated actin (supplementary Figures S2 and S3) and identified 1 % biotinylated lipids and 4 % biotinylated actin as suitable amounts which we used in the following experiments.

### Numerical Simulations

Theoretical predictions by Adeli Koudehi et al. have suggested that actin organization depends crucially on confinement and surface attachment.^46^ In order to explore the agreement of our experimental results with these simulations, we adopted their theoretical model. As such, we performed numerical simulations of interacting actin filaments under spherical confinement using Brownian dynamics (see Supplementary Methods).^46^ Semi-flexible actin filaments were modeled as beads connected by springs, with cross-linking represented by a short-range attraction with spring constant *k*_*atr*_. Polymerization from an initial number of filament seeds was simulated by addition of beads at one of the filament ends (representing the barbed end). The number of seeds was changed to achieve different final filament lengths. Boundary attraction was simulated as short-range attraction to the confining boundary. Simulated maximum intensity projections were performed as in Bidone et al.^47^

### Ring formation

Excitingly, and in accordance with the theoretical predictions, the most noticeable effect of membrane attachment was an increase in the formation of single actin rings. Although ring formation could still be observed without membrane-actin links, the introduction of membrane binding greatly enhances the probability of actin condensation into one single clearly discernible ring in vesicles. Membrane-bound actin rings have so far not been reported within synthetic vesicles.

Figure 2 highlights this effect of membrane attachment on the formation of actin rings. Figure 2A summarizes ring formation probabilities for three different bundlers, comparing conditions with and without membrane binding. We chose actin and bundler concentrations for which the formation of single rings is already relatively likely (30 % – 55 %) even without membrane attachment. In vesicles with membrane attached actin, probability of ring formation increased for all bundlers, and reaches up to 94% for actin bundled by vinculin and talin. While there is only a slight difference with and without membrane attachment for fascin, the likelihood of single ring occurrence increases by more than a factor of 1.7 for α-actinin and talin with vinculin (fluorescence image in Figure 2C).

**Figure 2:**
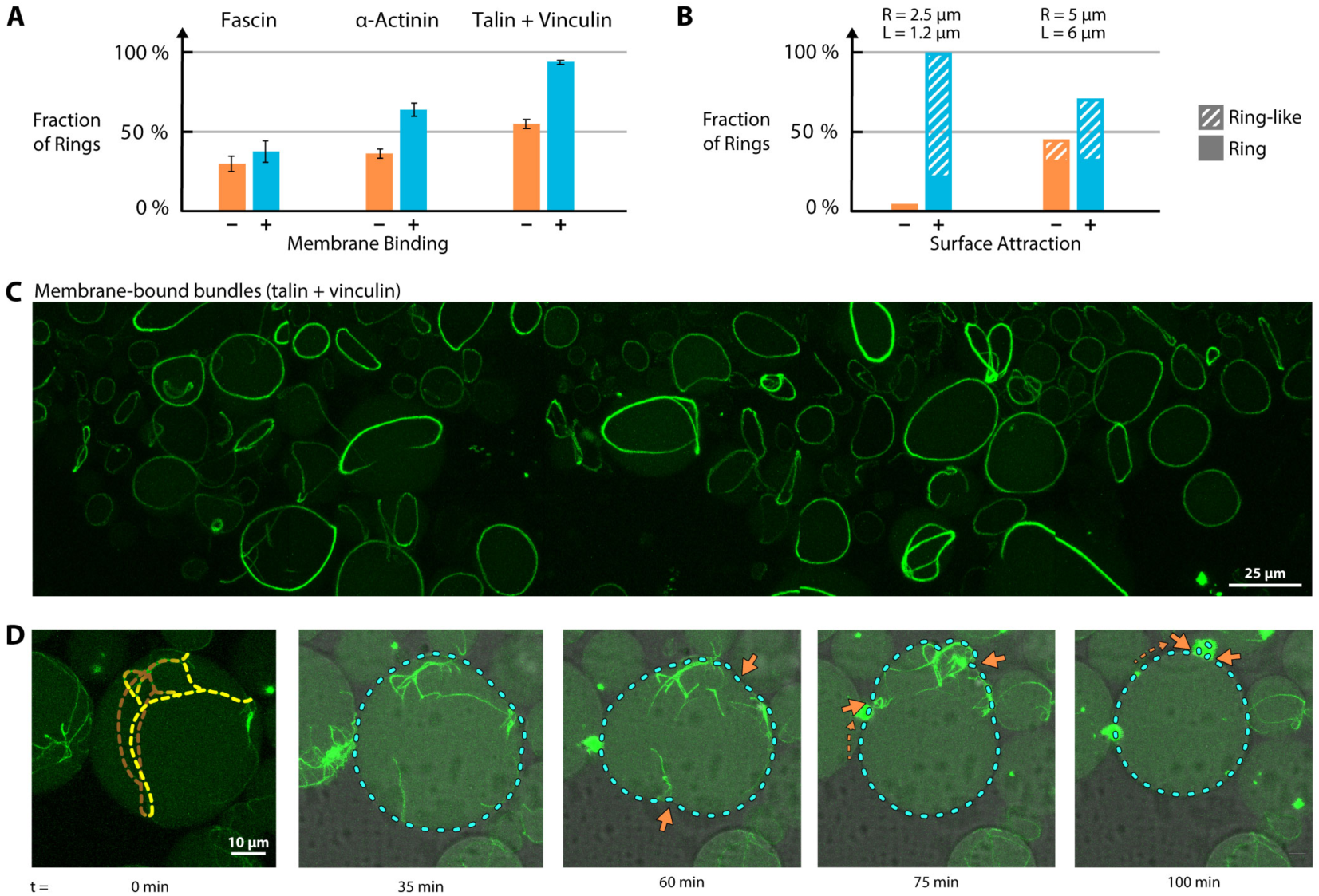
Formation of membrane-anchored actin rings. **A)** Membrane-binding promotes ring formation. Shown is the probability of the formation of single actin rings in GUVs (i.e. GUVs with one single unbranched actin bundle connected into a ring) in the absence and presence of membrane-anchoring. We use 2 µM actin in all cases, but due to differences in bundling activity different concentrations of bundling protein: 0.3 µM fascin, 1 µM α-actinin, 2 µM talin and 2 µM vinculin. **B)** Probability of ring formation for simulations with different initial parameters (*R*: Vesicle radius, *L*: Filament lengths). Snapshots from all simulation shown in supplementary Figure S5. **C)** Condition with particularly robust ring formation: actin bundled by talin with vinculin and bound to the membrane. Movie S3 shows a 3D view of this image. Supplementary Figure S4 shows rings formed by other bundling proteins. **D)** Time-series of a contracting ring-like structure in a GUV. Only the mid-section of the vesicle is shown, top and bottom are missing, but the yellow dotted lines in the first frame show the approximate position of the bundles. Orange arrows indicate membrane deformations (vesicle constriction). The partially visible vesicle on the left can be seen to undergo a similar transition from a large actin network (t = 0) to myosin-constricted cluster (t = 60 min).

In simulations, Adeli Koudehi et al. found that boundary attraction in the case of spherical confinement enhances the probability of ring formation from bundled filaments.^46^ However, the effect of boundary attraction in their work was studied for filament lengths larger than the confining diameter, whereas in experiments, we observed increased ring formation for vesicles and actin concentrations where the opposite should be true. We thus performed new simulations of actin filaments for concentrations chosen as in our experiments (*c* = 2 µM) and varied their lengths and confinement sizes (Figure 2B, supplementary Figure S5, Movie S4). We selected filament cross-linking simulation parameters that lead to bundle formation without filament sliding and associated bundle compaction *k*_*atr*_ = 2 pN/µm. Including surface boundary attraction greatly enhanced ring and ring-like structure formation for short filaments (length *L* = 1.2 µm) in small spheres (radius *R* = 2.5 µm). We also observed an enhancement of ring formation for filament lengths and sphere sizes comparable to that of our experiments (*L* = 6 µm, *R* = 5 µm), including when we increased the persistence length of individual actin filaments to simulate cross-linking induced bundle stiffening (Figure S5). Inspired by modeling results implying that the probability of ring formation depends on compartment size^46^, we analyzed our experimental data to confirm that rings preferably form in smaller vesicles (supplementary Figure S6).

The rings observed here can be assumed to mimic reorganization of actin which occurs during the last stages of cell division. In order to take this analogy one step further, we included muscle myosin II with the ultimate goal of forming a contractile actomyosin ring. Constriction of actin rings was shown before by Miyazaki et al., who demonstrated myosin-mediated contraction in a less cell-like system. They used actin bundled by depletion forces in water-in-oil droplets and showed that the behavior reproduced by this system, has a striking resemblance to constricting cell division rings.^48^

In our vesicles, the addition of myosin complicated the formation of single actin bundle rings. We used low concentrations of myosin II, such that the effect of myosin activity on bundling was minimized and motor-induced constriction slow enough to be observed while imaging. Although it appeared that the appropriate assay conditions for homogeneously contracting single rings have not yet been met in our giant vesicles, in few instances, we were able to observe the constriction of membrane-anchored ring-like structures along with membrane deformations. Figure 2D and Movie S5 show a time series of such a vesicle over the course of 2 hours. In accordance with our expectations, in this minimal system without further ring-stabilizing components, the constricting actomyosin ring eventually slides along the membrane and collapses into a single condensate on one side of the vesicle, a behavior that has been seen in yeast cells lacking cell walls.^36^ Such an arbitrary local collapse is not too surprising, as coordinated ring constriction in the cell is a highly spatially regulated process involving hundreds of proteins. Clearly, additional cellular machinery is required to stabilize the position of the ring, and membrane geometry and fluidity likely play additional roles. Figure 2D shows how the actomyosin ring initially deforms the vesicle membrane (orange arrows), leading to a furrow-like indentation. The entire timeseries without overlays is shown in Movie S5. Our experiments clearly show that the actin bundles are firmly attached to the inner leaflet of the vesicle membrane and that active forces are exerted by the motor proteins, capable of deforming the vesicle.

An additional effect of contraction of membrane-bound actomyosin is a change in the x-y cross section area of the vesicle after contraction occurs. This effect also appears for vesicles without initial furrow constriction. This is likely due to crumpling of the membrane into the actomyosin contraction point, which decreases membrane area (while vesicle volume is largely preserved) and increases membrane tension. As a result, vesicles that are initially slightly deflated become more spherical as a result of actomyosin contraction. Supplementary Figure S8 shows a DIC image of the actomyosin contraction point in Figure 2D and more examples of vesicles with shrinking x-y cross section.

### Membrane attachment shapes actin organization by curvature induction

We performed a series of experiments with the simplest bundling protein fascin to further investigate the effects of membrane binding on bundle morphology. We notice that membrane-binding primarily affects the curvature of the bundles: While actin with fascin forms very straight bundles that are just generally confined by the membrane, we see that membrane bound fascin bundles often adopt the exact curvature of the membrane (Figure 3A). Figure 3B shows a histogram for the distribution of bundle curvatures in these vesicles. The histogram shows a much broader distribution for unbound bundles, with a maximum at low curvatures, while the maximum for membrane-bound actin bundles is centered around the curvature of the membrane (relative curvature = 1.0).

**Figure 3:**
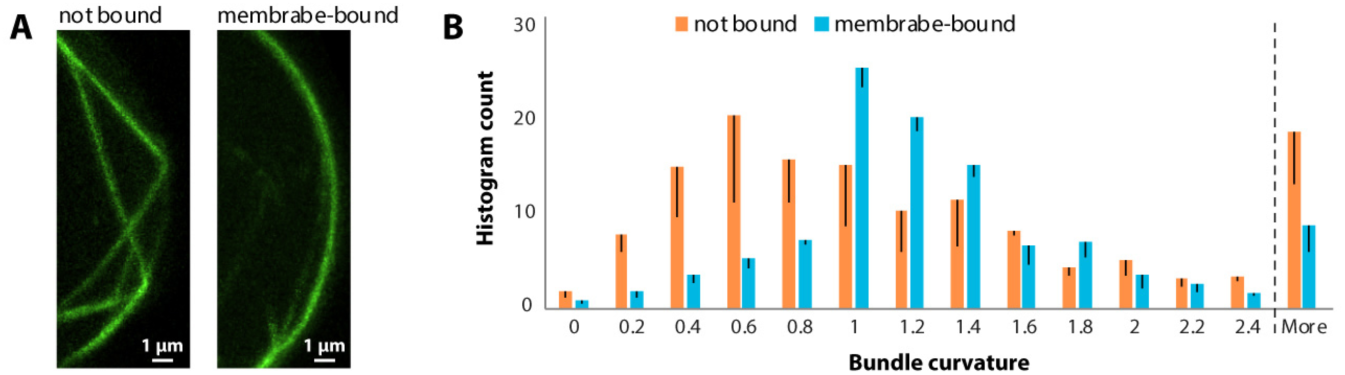
Membrane-attachment affects curvature of actin bundles. **A)** Each image shows a section of the membrane of a GUV with actin bundles bundled by fascin (2 µM actin, 0.2 µM fascin). Actin is in green. Unattached bundles have long sections with zero curvature, while membrane-bound bundles follow the curvature of the membrane. **B)** Distribution of curvatures for actin bundles in vesicles with and without membrane-binding. n = 5 for each condition. Confocal z-stacks were converted into 3D information (see Figure 1D). Bundles were then divided into small segments and their curvature was measured. Curvature is normalized by membrane curvature, so that a curvature of 1.0 equals the curvature of the membrane. More details about the analysis can be found in the supplements.

Despite this difference on a small scale, the general distribution of bundles within the vesicles seems to be largely independent on the presence of actin-membrane linkers. Figure 4A shows a set of conditions with and without membrane binding. We quantified the average actin distribution for each condition and find that (with few exceptions) actin is consistently positioned in close proximity to the membrane, with only minor differences between conditions with and without membrane linkers (Figure 4C and D). Our results indicate that if bundles are sufficiently long, their confinement by the vesicle boundary forces them to bend and concentrate at the inner surface. We note that for 2 µM actin (Figure 4A, top row) even low concentrations of fascin are sufficient to cause this effect, while we do observe that the thickness of the bundles increases with higher concentrations (supplementary Figure S9).

**Figure 4:**
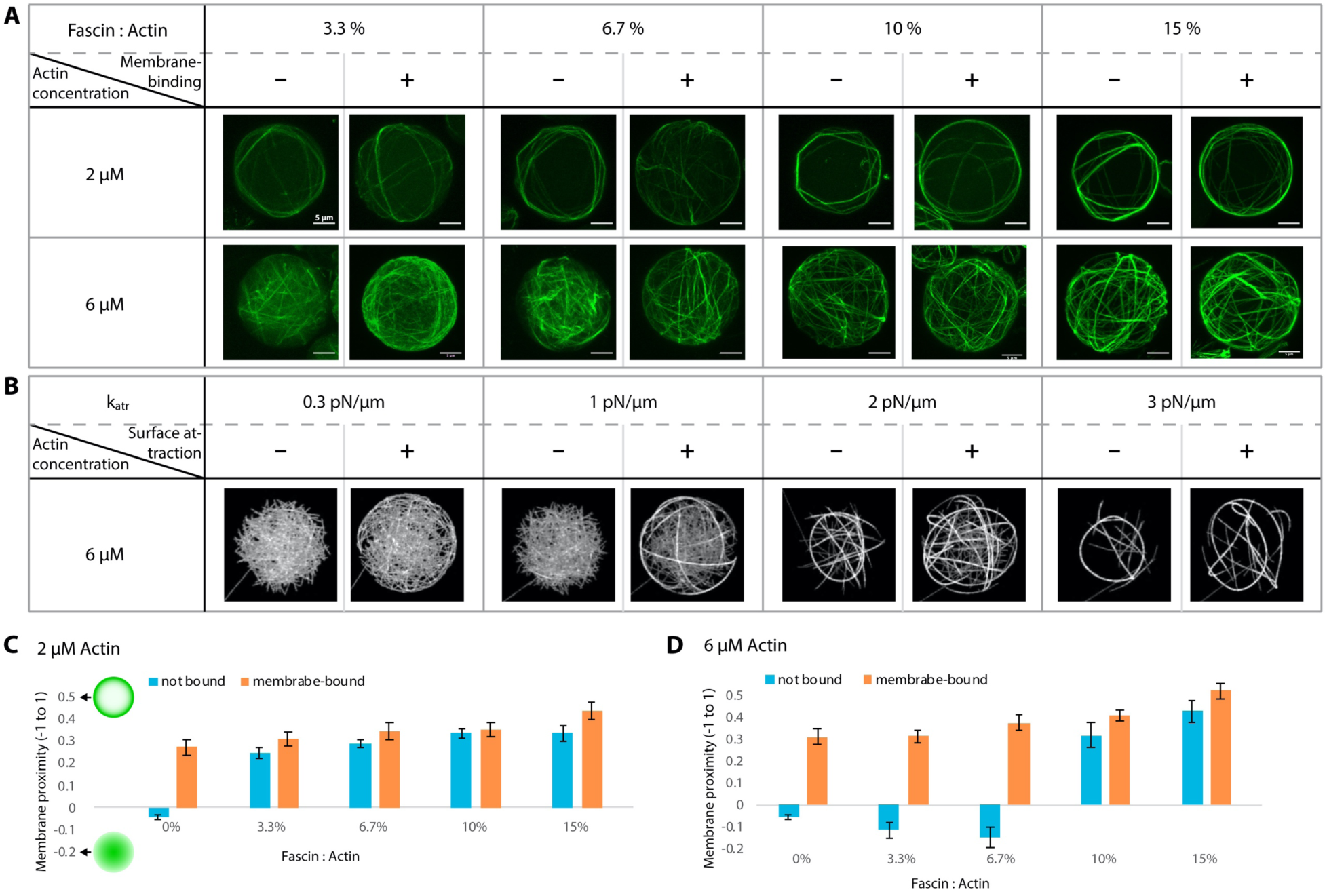
Actin organization in dependence on bundler to actin ratio and membrane binding. Actin is positioned close to the membrane regardless of membrane-anchoring. **A)** Overview of conditions with varying actin concentration (2 µM and 6 µM), fascin to actin ratios (3.3:100 to 15:100) and with and without actin-membrane binding. Supplemental Movie S6 shows a 3D version of this figure panel. **B)** Simulations for similar conditions as 6 µM conditions in (A). **C)** Average membrane proximity of actin signal for vesicles with 2 µM actin. Normalized range from −1 (all actin in the center of the vesicle) to +1 (all actin on the membrane). n = 10 for each condition. **D)** Membrane proximity of actin signal for vesicles with 6 µM actin. n = 10 for each condition.

At higher actin concentrations (6 µM) and low fascin to actin ratios (3.3% and 6.7%), bundles were shorter and thus more homogeneously distributed in the vesicles when not bound to the membrane (Figure 4D). Interestingly, we note that membrane-binding affects the threshold at which long actin bundles form: We observe long bundles at a fascin to actin ratio of 6.7% when we include membrane-linkers (see also supplementary Figure S10). These observations agree with corresponding simulations (Figure 4B).

In our experiments, ring-like structures consistently form at 2 µM actin, while at 6 µM actin, multiple bundles arrange themselves into cortex-like structures that do not condense into single rings. In our simulations, we see similar cortex-like morphologies in the early stages, but at longer times these condense into rings, both for low and high actin concentrations (supplementary Figure S11). Most likely this is a result of (1) the smaller confinement size we chose due to computation limitations, and (2) absence of a maximum bundle thickness, unlike in experiments.^49^

### Shaping the membrane compartment

Lipid membranes are highly flexible, and the shape of GUVs is mostly determined by the osmotic pressure inside the vesicle with respect to its enviroment. If this pressure is low, i.e. vesicles are osmotically deflated, strong deviations from the spherical shape are possible, and additional mechanical determinants, such as external forces or an encapsulated cytoskeleton induce arbitrary shapes of the vesicles,^31^ as can be seen in Figure 5A. The experiment we performed in Figure 5B further confirms the role of an artifcial cytoskeleton in determining vesicle shapes: By imaging deformed cytoskeletal vesicles with increased laser power on our confocal microscope, the actin filaments depolymerize after some time due to photo-damage^50^, relaxing the cytoskeleton-inferred shape determinants and leaving the deflated vesicles without internal support. This leads to dramatic changes in their shape, usually by taking on a spheroid (oblate) shape.

**Figure 5:**
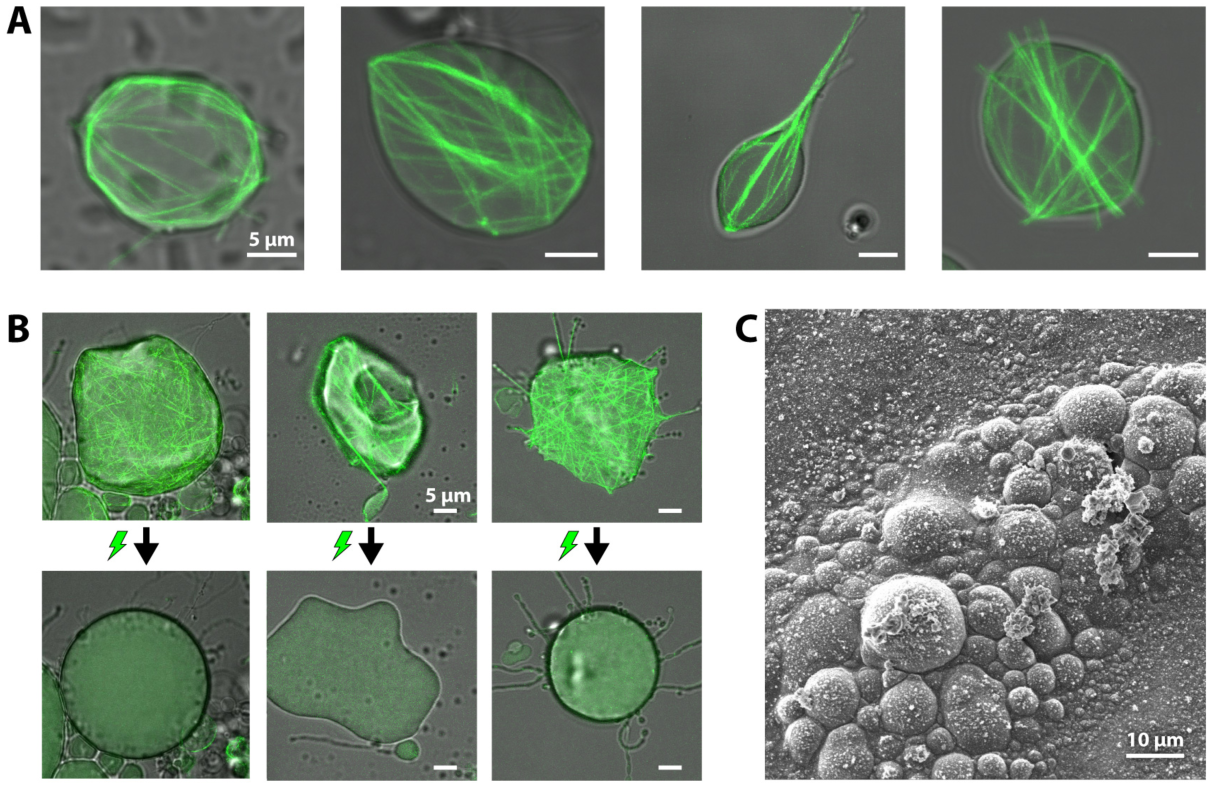
Actin cortex governs vesicle shape in osmotically deflated vesicles. **A)** Variety of vesicle shapes produced by different morphologies of encapsulated actin networks, some with stabilizing cortices and others with filopodia-like membrane protrusions. **B)** Upon exposure to photo damage through increased laser power, cytoskeletal vesicles lose their stabilizing actin cortex and take on a round shape. All three examples are shown in Movie S7. **C)** Vesicles with a stabilizing artificial actin cortex can be dried, frozen and imaged using cryo scanning electron microscopy.

These stabilizing cortices of actin bundles can even protect the vesicles, e.g., against the unfavorable conditions of sample preparation for cryo-electron microscopy, specifically the drying of the sample (removal of the surrounding aqueous phase): Figure 5C shows a cryo scanning electron microscopy image of frozen cytoskeletal vesicles. When we try to freeze and image vesicles without an encapsulated actin cortex or with actin bundles that are not attached to the membrane, vesicles rarely survive the process (Supplementary Figure S12). It has been shown in previous work that GUVs can be stabilized through a shell of cross-linked material on the membrane of GUVs, not only for unbundled actin^51^, but also with other proteins^52^, as well as DNA origami^53^. Here, we show that heterogeneously distributed, higher-order structures achieve a similar mechanical effect.

## Discussion

In this work, we succeeded in reconstituting defined ring-like actomyosin structures in giant unilamellar vesicles (GUVs). With respect to a suitable protein machinery that may serve as a minimal divisome for protocells, this constitutes the first important starting point for assembling contractile rings of sufficiently large sizes. To this end, we encapsulated a reaction mix into the vesicles that causes actin to polymerize, bundle, bind to the vesicle membrane and even contract. We show that the bundle networks can be highly organized and, under many conditions, reproducibly cross-linked into single rings.

Comparable ring formation has been shown by Miyazaki et al., who assembled actin into rings by depletion forces through the crowding agent methylcellulose, while confined in small water-in-oil droplets.^48^ These rings may contract, but due to the lack of surface attachment in this system, are unable to exert forces on the compartment interface. The formation of rings due to the extension of biopolymer bundles in confinement is known from both, theory^46,54^ and other experimental systems^30,55,56^. However, here we did not only form actin rings bundled by various physiological factors, but for the first time managed to attach these to the compartmentalizing lipid bilayer, such that ring contraction may result in dramatic transformation of the respective compartment from the inside.

We have shown, as a proof of principle, that myosin mediated constriction of such ring-like membrane-bound actin structures can be induced in lipid membrane vesicles. The vesicle deformations (Figure 2D) demonstrate the strength of the membrane anchoring. The final contracted state resembles myosin-induced symmetry breaking observed in other actomyosin *in vitro* systems under confinement.^18,26,57^ For example, Tsai et al. encapsulated a contractile actomyosin system in vesicles that condensed into large clusters.^58^

Further assay improvement and very likely, additional components will be necessary to accomplish a complete division of a cell-sized vesicle compartment. Unless membrane area can be expanded at the same time, the osmotic pressure inside the vesicle complicates a cell division-like symmetric constriction in the center of the vesicle, and in the absence of other geometric regulators, the fluidity of the membrane causes the ring-like bundles to “slip” and contract into a cluster in one location. We conclude that in order to achieve a binary fission through contraction of a single ring, more spatial determinants are required.

A behavior similar to what we observe, can be seen *in vivo* for the contraction of actomyosin rings in yeast protoplasts – yeast cells that were stripped from their cell walls.^36^ Stachowiak et al. beautifully demonstrated that in these spherical cells without cell walls, the contractile actomyosin ring slides along the cell membrane, collapsing into one point at the side of the cells. The absence of a cell wall in fission yeast results in both, a loss of their elongated shape, and in a lack of stabilizing the actomyosin ring in the cell center.^36^

By reconstituting a contractile actomyosin ring in giant unilamellar vesicles, we have made one essential step forward with regard to establishing a minimal system for active membrane vesicle division from the bottom up. Importantly, using this protein machinery from eukaryotes, large-size contractile ring structures could be generated that were for the first time attached to vesicle membranes from the inside. Our experiments demonstrate that while ring formation, membrane attachment, and contraction are not sufficient for division of these cell-like compartments, this system is ideal for identifying the additional parameters and components required. Further exploration will allow us to identify a minimal set of components that determine when, where, and how efficiently membrane compartments divide, providing vital insight into this key cellular process. In addition, the robust and reproducible methods used here provide a reliable platform for further reconstitution of the key processes of life beyond cell division.

## Acknowledgments

We thank Allen P. Liu for helpful discussions. We are grateful to Gunnar Goetz for experimental contributions, to David Rutkowski for contributing code to improve simulation efficiency and Julia Skrapits for help with simulations. We thank Giovanni Cardone and Martin Spitaler from the MPI-B Imaging Facility for assistance with data analysis. This work is part of the MaxSynBio consortium, which is jointly funded by the Federal Ministry of Education and Research of Germany and the Max Planck Society. CK is a recipient of the Humboldt Research Fellowship for Postdoctoral Researchers. NM acknowledges the Boehringer Ingelheim Foundation Plus 3 Program, and the European Research Council (ERC-CoG, 724209). DV, DH and MAK were supported by National Institutes of Health Grant R01GM114201. Use of the high-performance computing capabilities of the Extreme Science and Engineering Discovery Environment (XSEDE), which is supported by the National Science Foundation, project no. TG-MCB180021 is also gratefully acknowledged.

## References

1 Rottner, K., Faix, J., Bogdan, S., Linder, S. & Kerkhoff, E. Actin assembly mechanisms at a glance. J Cell Sci 130, 3427–3435, doi: 10.1242/jcs.206433 (2017).

2 Blanchoin, L., Boujemaa-Paterski, R., Sykes, C. & Plastino, J. Actin dynamics, architecture, and mechanics in cell motility. Physiol Rev 94, 235–263, doi: 10.1152/physrev.00018.2013 (2014).

3 Schwayer, C., Sikora, M., Slovakova, J., Kardos, R. & Heisenberg, C. P. Actin Rings of Power. Dev Cell 37, 493–506, doi: 10.1016/j.devcel.2016.05.024 (2016).

4 Doherty, G. J. & McMahon, H. T. Mediation, modulation, and consequences of membrane-cytoskeleton interactions. Annu Rev Biophys 37, 65–95, doi: 10.1146/annurev.biophys.37.032807.125912 (2008).

5 Bezanilla, M., Gladfelter, A. S., Kovar, D. R. & Lee, W.-L. Cytoskeletal dynamics: A view from the membrane. Journal of Cell Biology 209, 329–337, doi: 10.1083/jcb.201502062 (2015).

6 Liu, A. P. et al. Membrane-induced bundling of actin filaments. Nature Physics 4, 789–793, doi: 10.1038/nphys1071 (2008).

7 Ganzinger, K. A. & Schwille, P. More from less – bottom-up reconstitution of cell biology. J Cell Sci 132, doi: 10.1242/jcs.227488 (2019).

8 Liu, A. P. & Fletcher, D. A. Biology under construction: in vitro reconstitution of cellular function. Nat Rev Mol Cell Biol 10, 644–650, doi: 10.1038/nrm2746 (2009).

9 Kuhne, W. Untersuchungen uber das Protoplasma und die Contractilitat. (W. Engelmann, 1864).

10 Straub, F. Studies of the Inst. of Med. Chem. Univ. Szeged edit., by Szent-Györgyi, vol. II. Basel and New York: S. Karger, 3 (1942).

11 Szent-Györgyi, A. G. The Early History of the Biochemistry of Muscle Contraction. The Journal of General Physiology 123, 631–641, doi: 10.1085/jgp.200409091 (2004).

12 Mullins, R. D. & Hansen, S. D. In vitro studies of actin filament and network dynamics. Curr Opin Cell Biol 25, 6–13, doi: 10.1016/j.ceb.2012.11.007 (2013).

13 Smith, B. A., Gelles, J. & Goode, B. L. Single-molecule studies of actin assembly and disassembly factors. Methods Enzymol 540, 95–117, doi: 10.1016/B978-0-12-397924-7.00006-6 (2014).

14 Pollard, T. D. Actin and Actin-Binding Proteins. Cold Spring Harb Perspect Biol 8, doi: 10.1101/cshperspect.a018226 (2016).

15 Dogterom, M. & Koenderink, G. H. Actin-microtubule crosstalk in cell biology. Nat Rev Mol Cell Biol 20, 38–54, doi: 10.1038/s41580-018-0067-1 (2019).

16 Carvalho, K. et al. Cell-sized liposomes reveal how actomyosin cortical tension drives shape change. Proc Natl Acad Sci U S A 110, 16456–16461, doi: 10.1073/pnas.1221524110 (2013).

17 Caorsi, V. et al. Cell-sized liposome doublets reveal active tension build-up driven by acto-myosin dynamics. Soft Matter 12, 6223–6231, doi: 10.1039/c6sm00856a (2016).

18 Carvalho, K. et al. Actin polymerization or myosin contraction: two ways to build up cortical tension for symmetry breaking. Philos Trans R Soc Lond B Biol Sci 368, 20130005, doi: 10.1098/rstb.2013.0005 (2013).

19 Vogel, S. K., Petrasek, Z., Heinemann, F. & Schwille, P. Myosin motors fragment and compact membrane-bound actin filaments. Elife 2, e00116, doi: 10.7554/eLife.00116 (2013).

20 Sonal et al. Myosin-II activity generates a dynamic steady state with continuous actin turnover in a minimal actin cortex. J Cell Sci 132, doi: 10.1242/jcs.219899 (2018).

21 Murrell, M. & Gardel, M. L. Actomyosin sliding is attenuated in contractile biomimetic cortices. Mol Biol Cell 25, 1845–1853, doi: 10.1091/mbc.E13-08-0450 (2014).

22 Gopfrich, K., Platzman, I. & Spatz, J. P. Mastering Complexity: Towards Bottom-up Construction of Multifunctional Eukaryotic Synthetic Cells. Trends Biotechnol 36, 938–951, doi: 10.1016/j.tibtech.2018.03.008 (2018).

23 Schwille, P. et al. MaxSynBio: Avenues Towards Creating Cells from the Bottom Up. Angew Chem Int Ed Engl 57, 13382–13392, doi: 10.1002/anie.201802288 (2018).

24 Mulla, Y., Aufderhorst-Roberts, A. & Koenderink, G. H. Shaping up synthetic cells. Physical Biology 15, 041001, doi: 10.1088/1478-3975/aab923 (2018).

25 Bashirzadeh, Y. & Liu, A. P. Encapsulation of the cytoskeleton: towards mimicking the mechanics of a cell. Soft Matter, doi: 10.1039/c9sm01669d (2019).

26 Loiseau, E. et al. Shape remodeling and blebbing of active cytoskeletal vesicles. Sci Adv 2, e1500465, doi: 10.1126/sciadv.1500465 (2016).

27 Durre, K. et al. Capping protein-controlled actin polymerization shapes lipid membranes. Nat Commun 9, 1630, doi: 10.1038/s41467-018-03918-1 (2018).

28 Maan, R., Loiseau, E. & Bausch, A. R. Adhesion of Active Cytoskeletal Vesicles. Biophys J 115, 2395–2402, doi: 10.1016/j.bpj.2018.10.013 (2018).

29 Honda, M., Takiguchi, K., Ishikawa, S. & Hotani, H. Morphogenesis of liposomes encapsulating actin depends on the type of actin-crosslinking. J Mol Biol 287, 293–300, doi: 10.1006/jmbi.1999.2592 (1999).

30 Limozin, L. & Sackmann, E. Polymorphism of Cross-Linked Actin Networks in Giant Vesicles. Physical Review Letters 89, doi: 10.1103/PhysRevLett.89.168103 (2002).

31 sai, F. C. & Koenderink, G. H. Shape control of lipid bilayer membranes by confined actin bundles. Soft Matter 11, 8834–8847, doi: 10.1039/c5sm01583a (2015).

32 Majumder, S., Wubshet, N. & Liu, A. P. Encapsulation of complex solutions using droplet microfluidics towards the synthesis of artificial cells. Journal of Micromechanics and Microengineering 29, doi: 10.1088/1361-6439/ab2377 (2019).

33 Sato, Y. & Takinoue, M. Creation of Artificial Cell-Like Structures Promoted by Microfluidics Technologies. Micromachines 10, doi: 10.3390/mi10040216 (2019).

34 Supramaniam, P., Ces, O. & Salehi-Reyhani, A. Microfluidics for Artificial Life: Techniques for Bottom-Up Synthetic Biology. Micromachines 10, doi: 10.3390/mi10050299 (2019).

35 Robinson, T. Microfluidic Handling and Analysis of Giant Vesicles for Use as Artificial Cells: A Review. Advanced Biosystems 3, doi: 10.1002/adbi.201800318 (2019).

36 Stachowiak, M. R. et al. Mechanism of cytokinetic contractile ring constriction in fission yeast. Dev Cell 29, 547–561, doi: 10.1016/j.devcel.2014.04.021 (2014).

37 Hürtgen, D., Härtel, T., Murray, S. M., Sourjik, V. & Schwille, P. Functional Modules of Minimal Cell Division for Synthetic Biology. Advanced Biosystems 3, 1800315, doi: 10.1002/adbi.201800315 (2019).

38 Kretschmer, S., Ganzinger, K. A., Franquelim, H. G. & Schwille, P. Synthetic cell division via membrane-transforming molecular assemblies. BMC Biol 17, 43, doi: 10.1186/s12915-019-0665-1 (2019).

39 Exterkate, M. & Driessen, A. J. M. Synthetic Minimal Cell: Self-Reproduction of the Boundary Layer. ACS Omega 4, 5293–5303, doi: 10.1021/acsomega.8b02955 (2019).

40 Schwille, P. Division in synthetic cells. Emerging Topics in Life Sciences, doi: 10.1042/etls20190023 (2019).

41 Abkarian, M., Loiseau, E. & Massiera, G. Continuous droplet interface crossing encapsulation (cDICE) for high throughput monodisperse vesicle design. Soft Matter 7, 4610–4614, doi: 10.1039/C1SM05239J (2011).

42 Litschel, T., Ramm, B., Maas, R., Heymann, M. & Schwille, P. Beating Vesicles: Encapsulated Protein Oscillations Cause Dynamic Membrane Deformations. Angew Chem Int Ed Engl 57, 16286–16290, doi: 10.1002/anie.201808750 (2018).

43 Jansen, S. et al. Mechanism of actin filament bundling by fascin. J Biol Chem 286, 30087–30096, doi: 10.1074/jbc.M111.251439 (2011).

44 Winkelman, J. D. et al. Fascin- and alpha-Actinin-Bundled Networks Contain Intrinsic Structural Features that Drive Protein Sorting. Curr Biol 26, 2697–2706, doi: 10.1016/j.cub.2016.07.080 (2016).

45 Hüttelmaier, S. et al. Characterization of the actin binding properties of the vasodilator-stimulated phosphoprotein VASP. FEBS Letters 451, 68–74, doi: 10.1016/s0014-5793(99)00546-3 (1999).

46 Adeli Koudehi, M., Rutkowski, D. M. & Vavylonis, D. Organization of Associating or Crosslinked Actin Filaments in Confinement. Cytoskeleton 76, 532–548, doi: 10.1002/cm.21565 (2019).

47 Bidone, T. C., Tang, H. & Vavylonis, D. Dynamic network morphology and tension buildup in a 3D model of cytokinetic ring assembly. Biophys J 107, 2618–2628, doi: 10.1016/j.bpj.2014.10.034 (2014).

48 Miyazaki, M., Chiba, M., Eguchi, H., Ohki, T. & Ishiwata, S. Cell-sized spherical confinement induces the spontaneous formation of contractile actomyosin rings in vitro. Nat Cell Biol 17, 480–489, doi: 10.1038/ncb3142 (2015).

49 Claessens, M. M., Semmrich, C., Ramos, L. & Bausch, A. R. Helical twist controls the thickness of F-actin bundles. Proc Natl Acad Sci U S A 105, 8819–8822, doi: 10.1073/pnas.0711149105 (2008).

50 van der Gucht, J., Paluch, E., Plastino, J. & Sykes, C. Stress release drives symmetry breaking for actin-based movement. Proc Natl Acad Sci U S A 102, 7847–7852, doi: 10.1073/pnas.0502121102 (2005).

51 Schafer, E., Vache, M., Kliesch, T. T. & Janshoff, A. Mechanical response of adherent giant liposomes to indentation with a conical AFM-tip. Soft Matter 11, 4487–4495, doi: 10.1039/c5sm00191a (2015).

52 Dieluweit, S. et al. Mechanical properties of bare and protein-coated giant unilamellar phospholipid vesicles. A comparative study of micropipet aspiration and atomic force microscopy. Langmuir 26, 11041–11049, doi: 10.1021/la1005242 (2010).

53 Kurokawa, C. et al. DNA cytoskeleton for stabilizing artificial cells. Proc Natl Acad Sci U S A 114, 7228–7233, doi: 10.1073/pnas.1702208114 (2017).

54 Vetter, R., Wittel, F. K. & Herrmann, H. J. Morphogenesis of filaments growing in flexible confinements. Nat Commun 5, 4437, doi: 10.1038/ncomms5437 (2014).

55 Keber, F. C. et al. Topology and dynamics of active nematic vesicles. Science 345, 1135–1139, doi: 10.1126/science.1254784 (2014).

56 Baumann, H. & Surrey, T. Motor-mediated cortical versus astral microtubule organization in lipid-monolayered droplets. J Biol Chem 289, 22524–22535, doi: 10.1074/jbc.M114.582015 (2014).

57 Ierushalmi, N. et al. Centering and symmetry breaking in confined contracting actomyosin networks. eLife 9, e55368, doi: 10.7554/eLife.55368 (2020).

58 Tsai, F.-C., Stuhrmann, B. & Koenderink, G. H. Encapsulation of Active Cytoskeletal Protein Networks in Cell-Sized Liposomes. Langmuir 27, 10061–10071, doi: 10.1021/la201604z (2011).

